# Category-based attention facilitates memory search

**DOI:** 10.1101/2023.12.08.570779

**Authors:** Linlin Shang, Lu-Chun Yeh, Yuanfang Zhao, Iris Wiegand, Marius V. Peelen

**Author notes:** Corresponding authors: Radboud University 6525 GD Nijmegen The Netherlands.

## Abstract

We often need to decide whether the object we look at is also the object we look for. When we look for one specific object, this process can be facilitated by preparatory feature-based attention. However, when we look for multiple objects at the same time (e.g., the products on our shopping list) such a strategy may no longer be possible, as research has shown that we can actively prepare to detect only one object at a time. Therefore, looking for multiple objects may additionally involve search in long-term memory, slowing down decision making. Interestingly, however, previous research has shown that memory search can be very efficient when distractor objects are from a different category than the items in the memory set. Here, using EEG, we show that this efficiency is supported by top-down attention at the category level. In Experiment 1, human participants (both sexes) performed a memory search task on individually presented objects of the same or different category as the objects in the memory set. We observed category-level attentional modulation of distractor processing from ∼150 ms after stimulus onset, expressed both as an evoked response modulation and as an increase in decoding accuracy of same-category distractors. In Experiment 2, memory search was performed on two concurrently presented objects. When both objects were distractors, spatial attention (indexed by the N2pc component) was directed to the object that was of the same category as the objects in the memory set. Together, these results demonstrate how attention can facilitate memory search.

**Significance statement:** When we are in the supermarket, we repeatedly decide whether a product we look at (e.g., a banana) is on our memorized shopping list (e.g., apples, oranges, kiwis). This requires searching our memory, which takes time. However, when the product is of an entirely different category (e.g., dairy instead of fruit), the decision can be made quickly. Here, we used EEG to show that this between-category advantage in memory search tasks is supported by top-down attentional modulation of visual processing: The visual response evoked by distractor objects was modulated by category membership, and spatial attention was quickly directed to the location of within-category (vs. between-category) distractors. These results demonstrate a close link between attention and memory.

## Introduction

Visual object processing is modulated by top-down goals. For example, the same object evokes a stronger neural response in visual cortex when the object is a target as compared to when it is a distractor (e.g., Chelazzi et al., 1993; Bansal et al., 2014). These modulations have typically been studied in the context of attention, with a top-down attentional set (or “template”) modulating visual processing (Desimone & Duncan, 1995). Such templates can operate at different levels of the visual hierarchy, from simple visual features to high-level object categories (Battistoni et al., 2017).

While the mechanisms behind single-target detection have been extensively studied, much less is known about multiple-target detection (Ort & Olivers 2020), even though this task is common in daily life. For example, when we are in the supermarket, we must decide whether the product we look at is one of the (possibly many) products on our memorized shopping list. This task is typically referred to as a memory search task (Sternberg, 1966), as it involves searching memory for the currently fixated item. Indeed, as memory set size (MSS) increases, responses systematically slow down, reflecting the memory search process (Wolfe, 2012).

An important difference between single-target and multiple-target detection is that observers can no longer use an attentional template-based strategy when looking for multiple targets. This is because only one or two attentional templates can be activated at a given time (Houtkamp & Roelfsema, 2006; Olivers et al., 2011; Ort & Olivers 2020; van Moorselaar et al., 2014; Wolfe 2021). Accordingly, the attentional template-based modulation of visual object processing, as observed for single-target detection, may be absent for multiple-target detection (i.e., memory search). This is in line with findings from memory research, showing relatively late (∼300-500 msec) electroencephalography (EEG) responses over mid-frontal and parietal electrodes reflecting recognition memory (see Rugg & Curran, 2007 for a review), rather than the earlier (150-200 ms) attentional modulation observed in single-target detection tasks (VanRullen & Thorpe, 2001; Kaiser et al., 2016).

Interestingly, however, memory search efficiency depends on the categorical relationship between the objects in the memory set and the probe (Cunningham & Wolfe, 2012, 2014; Drew & Wolfe, 2014). If the probe (e.g., a banana) is of the same category as the items in the memory set (e.g., apple, pear, orange), search is inefficient, such that RT increases strongly with increasing MSS. However, when the probe is of a different category (e.g., an animal), search is highly efficient, such that RT increases only weakly with increasing MSS (Cunningham & Wolfe, 2012, 2014; Drew & Wolfe, 2014). These findings could reflect differences in memory search efficiency, such that between-category distractors are rejected efficiently because they are represented distinctly in long-term memory (Figure 1A). Alternatively, however, participants could use the shared category of the memory items to form a category-level attentional template, thereby efficiently rejecting between-category distractors even before commencing search in long-term memory (Cunningham & Wolfe, 2014; Figure 1B).

**Figure 1.**
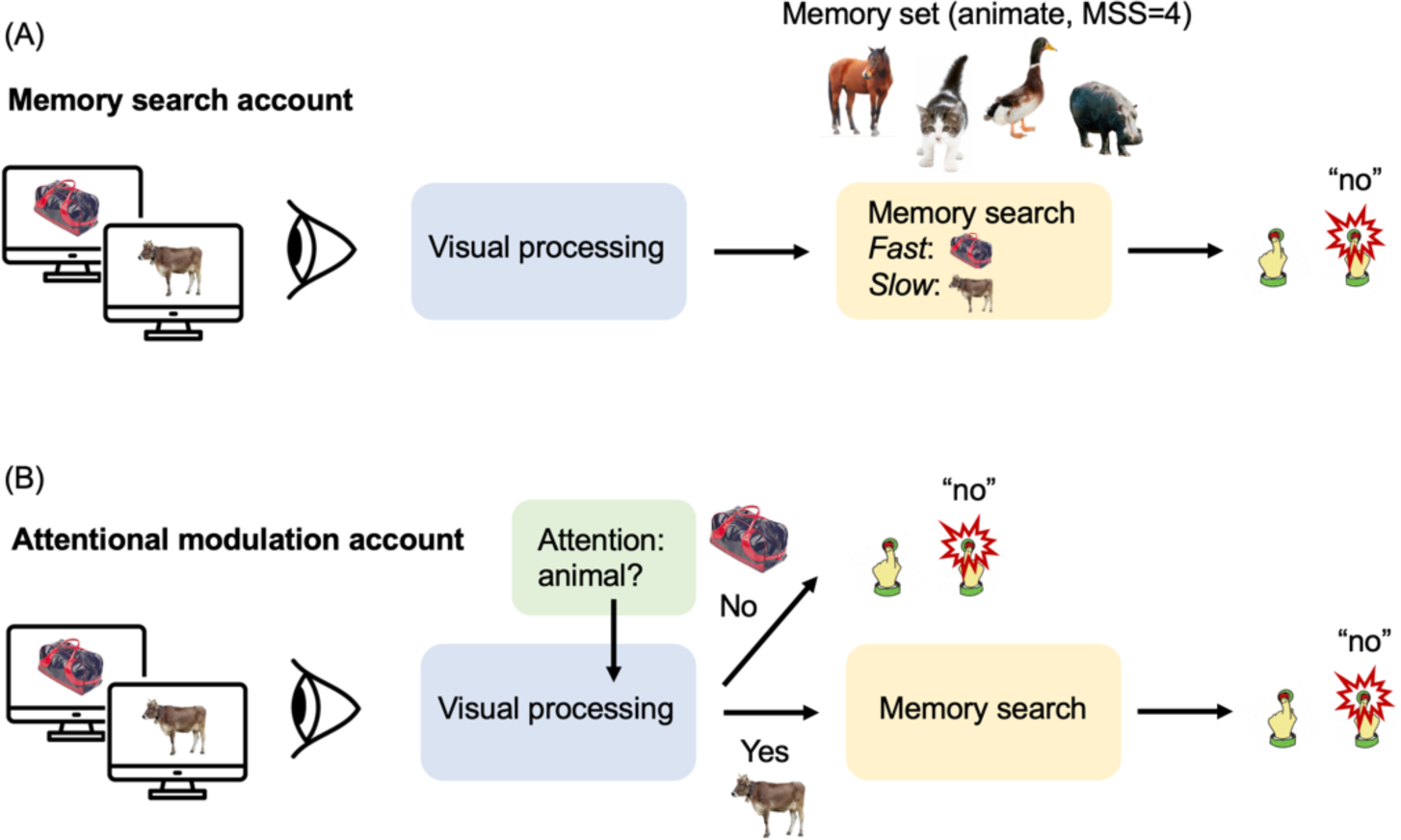
Schematic illustration of two accounts explaining category effects in memory search. The memory set in this example consists of four animals. (A) On a memory search account, all objects enter the memory search phase after visual object processing is completed. Faster RTs for between-category than within-category distractors are explained by differences in the efficiency of the memory search process. (B) On an attentional modulation account, participants attend to the category of the items in the memory set, modulating visual processing and resulting in the quick rejection of between-category distractors, largely avoiding memory search for these items. The current experiments were aimed at providing evidence for category-level attentional modulation of visual processing.

Here, in two experiments, we used EEG to test whether participants spontaneously form category-level attentional templates in a memory search task. We tested if, and when, visual object processing is modulated by category similarity in a memory search task. In Experiment 1, we found that distractors that were of the same category as the memorized items (within-category distractors) received more attention, and were therefore processed more strongly, than distractors that were of a different category (between-category distractors). In Experiment 2, we used two-object displays to show that this modulation results in the rapid allocation of spatial attention towards within-category distractors. Altogether, our results show that the between-category advantage in memory search is partly explained by category-level attentional modulation.

## Materials and Methods

### Participants

#### Experiment 1a

Forty-one participants (15 females; mean age, 23.81 years; age range, 20-35 years) were recruited from the online platform Prolific to arrive at a final sample size of 40. Twenty participants (8 females; mean age, 22.95 years; age range, 20-28 years) were assigned to the animate-category group, while 21 participants (7 females; mean age, 24.94 years; age range, 20-35 years) were assigned to the inanimate-category group. One participant in this group had to be excluded because of low accuracy (ACC) for set size 16 (<65%). All participants signed an online informed consent form and received 6 euro per hour for their participation in the experiment, which was approved by the Ethics Committee of the Faculty of Social Sciences, Radboud University Nijmegen.

#### Experiment 1b

We employed pwr [R] package to compute the sample size at the significance level of 0.05 with Cohen’s d 0.5 and power 0.8. Thirty-four participants were needed. We included 32 participants (23 females; age range, 18-30 years with *Mean* = 22.031 years and *SD* = 3.036) in Experiment 1b because of the Latin square design (see Experimental Design and Procedure). All participants had normal or corrected-to-normal vision. The participants gave written informed consent and received a gift card of 10 euro per hour for their participation. The study was approved by the Ethics Committee of the Faculty of Social Sciences, Radboud University Nijmegen.

#### Experiment 2

To arrive at 32 participants, as in Experiment 1b, 35 right-handed participants (11 females; age ranges from 18 to 35 years with *Mean* = 22.51 years and *SD* = 3.61) were recruited. Three participants were excluded due to missing more than 20% of trials after incorrect responses exclusion and artifact rejection. All participants had normal or corrected-to-normal vision. The participants gave written informed consent and received a gift card of 10 euro per hour for their participation. The study was approved by the Ethics Committee of the Faculty of Social Sciences, Radboud University Nijmegen.

### Stimuli

#### Experiment 1a

Stimuli consisted of full-color photographs of isolated animals (from Google Images) and inanimate objects (from Brady et al., 2008). Both categories had 30 subcategories (e.g., horse, cat, binoculars, bowl), and each subcategory consisted of 17 exemplar images, for a total of 1020 unique images. Stimulus size was 500 × 500 pixels. The experiment was programmed with PsychoPy v2020.2.3 (Peirce et al., 2019) and was hosted on Pavlovia.

#### Experiment 1b

The stimuli were the same as in Experiment 1a. The experiment was programmed with PsychoPy v2022.2.4 (Peirce et al., 2019) and ran on a 24-inch monitor (BenQ XL2420Z) with a refresh rate of 120 Hz and a resolution of 1920×1080. Participants were required to keep a distance of approximately 57 cm from the screen, and the stimuli subtended a visual angle of 4.9°.

#### Experiment 2

The stimuli were a subset of those used in Experiment 1, removing one subcategory of each superordinate category for a total of 986 full-color images of unique isolated objects. Stimuli were presented on a white background with a visual angle of 4°. The experiment was programmed with PsychoPy v2022.2.5 (Peirce et al., 2019) and was presented on a 24-inch monitor (BenQ XL2420Z) with a refresh rate of 120 Hz and a resolution of 1920×1080.

### Experimental Design and Procedure

#### Experiment 1a

The experimental design followed a 2 (animate/inanimate category group; between-subjects) × 2 (within/between category; within-subjects) × 5 (MSS 1/2/4/8/16; within-subjects) mixed factorial design. Category group was manipulated between subjects, such that each participant remembered either animate or inanimate objects in all blocks of the experiment. MSS was blocked, with one block for each MSS, for a total of 5 blocks per participant. Block order was randomized. Each block consisted of the following three phases (Figure 2):

(1) In the first phase (memorization), participants memorized the objects belonging to the memory set in that block. Object images were randomly assigned to the memory set, with the constraint that each image in the set came from a different subordinate category (e.g., horse, cat). All images in the memory set were new to the participant. In each trial, a central fixation cross appeared for 800 msec as a prompt, followed by an object presented in the center of the screen on a white background, one at a time, for 3000 msec with an inter-stimulus interval (ISI) of 950 msec. Participants were instructed to memorize the objects without giving a response.

(2) In the second phase (memory test), participants again viewed the objects but now had to indicate, with a button press, whether the object belonged to the memory set (press “z”) or not (press “m”). This task was self-paced. Non-target objects were randomly drawn from the same sub-ordinate categories as the target objects (e.g., a different cat). Half of the objects were targets and half were not, presented in random order without repetition. Participants had to be at least 80% correct on two subsequent tests to be able to proceed to the next phase of the experiment. If they did not meet this criterion, they would repeat Phase 1.

(3) In the third and main phase (memory search) of each block, participants performed a speeded old/new recognition task, deciding for each object whether or not it was part of the memory set. This phase consisted of 60 trials, presented in random order. Twenty percent of the trials (12 trials) showed an image selected from the memory set, while the remaining 48 trials were target-absent trials. Of these target-absent trials, 24 belonged to the animate category and 24 belonged to the inanimate category. Therefore, depending on the category group the participant was assigned to, these could be either within- or between-category distractors (relative to the memory set). In each trial, a central fixation cross appeared for 800 msec as a prompt, followed by an object presented in the center of the screen on a white background for 200 msec with an ISI of 1800 msec. Participants had to indicate whether the object belonged to the memory set within 2000 msec. The target images were randomly drawn from the memory set, such that these could repeat within a block. However, all the non-target images were unique across the whole experiment.

**Figure 2.**
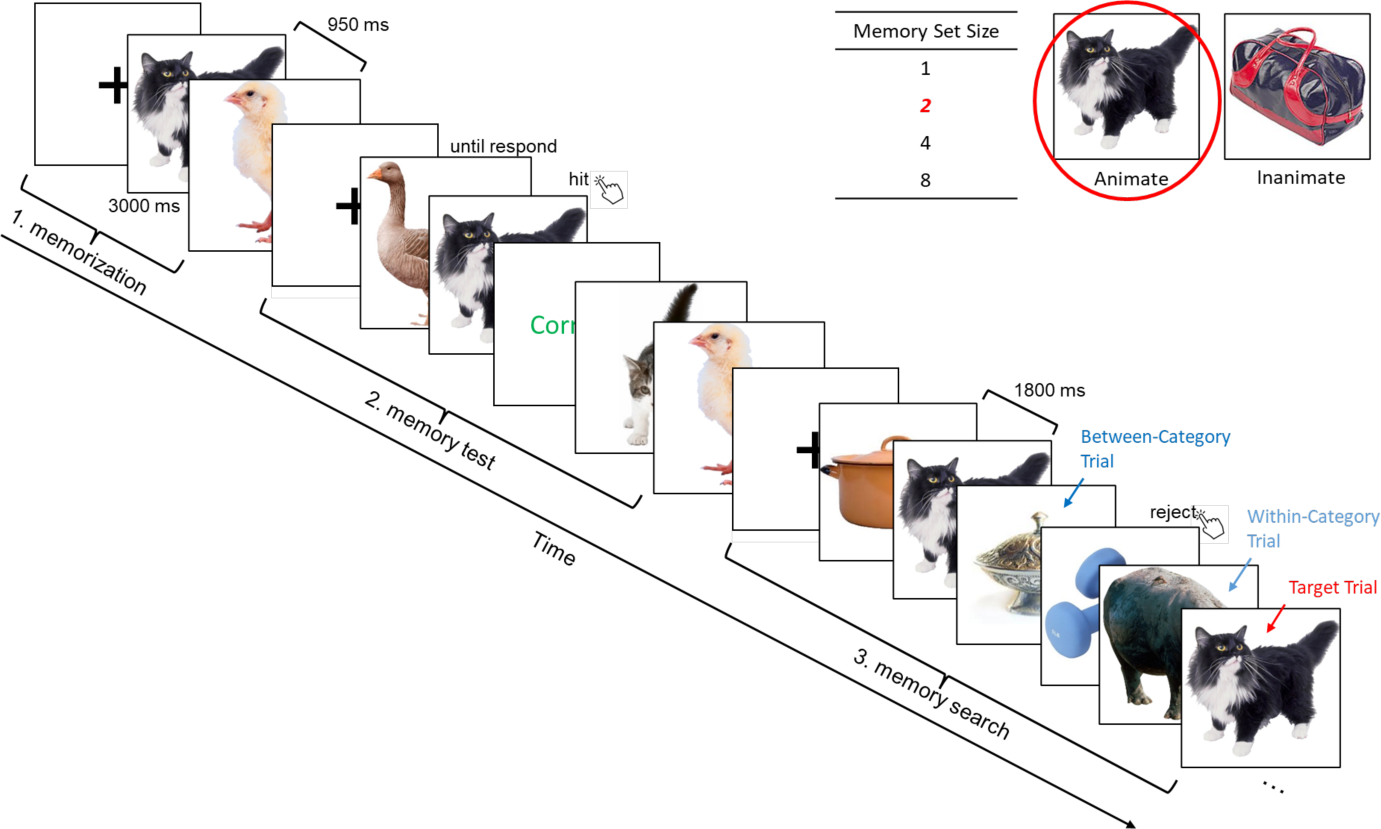
Illustration of the three phases in each block of Experiments 1a and b. The example presents an animate block with two targets (i.e., MSS 2).

#### Experiment 1b

##### Memory Search

The memory search design generally followed the design of Experiment 1a with minor adjustments. To simplify the experimental procedure, set size 16 was removed. Furthermore, unlike Experiment 1a, the category (animate or inanimate) was manipulated within participants (i.e., participants memorized the targets from both animate and inanimate categories). As in Experiment 1a, each MSS was designed as a block; there were therefore 8 blocks in total in the memory task. A Latin square design was used to order the four MSS blocks, with category order within MSS randomly determined. Unlike Experiment 1a, the fixation cross was always presented except during the stimulus presentation, and the ISI in the memory search phase was jittered between 1800 and 2300 msec. Other procedures were identical to Experiment 1a (Figure 2).

##### Visual Oddball Task

A visual oddball task was included to measure visually evoked response patterns to animate and inanimate objects without a memory search task. These data were used to train an animate/inanimate classifier, ensuring that classifier training was done on independent data. In each run, 50 animate and 50 inanimate objects were shown for 200 msec, one by one, with an ISI of 1800-2300 msec, in random order. The 100 objects were randomly selected from the same stimulus pool as used for the memory search task. Objects that were selected for the oddball task were not selected for the memory search task. In addition to the objects, there were ten two-digit numbers that were randomly interspersed. Participants pressed a button when seeing one of these numbers. In total, participants performed the oddball task three times. These runs were preceded and followed by two memory search blocks.

##### EEG Acquisition and Pre-Processing

Scalp EEG signals were recorded with a customized 64-channel active electrode actiCAP system with 500 Hz sampling rate. AFz served as ground electrode, and TP9 placed on left mastoid as a reference electrode. FT9/FT10 and Fp1/Fp2 were reset to left/right and up/down eye movement recorder. Impedance of all the electrodes was kept below 20 kΩ. The EEG data were pre-processed in Python 3.10 using custom code adapted from MNE toolbox (Gramfort, 2013). All the data were bandpass filtered (0.1 and 40 Hz) and resampled to 250 Hz. Each trial epoch was segmented from -200 to 800 msec relative to the onset of the object. Only epochs with correct responses were included in further analyses. Then, independent component analysis (ICA) was performed for each subject to remove components of eye movements and blinks. Finally, the ICA-corrected data were re-referenced to the average of all channels, and were baseline corrected by subtracting the mean activity from -200 to 0 msec.

#### Experiment 2

As in Experiment 1b, Experiment 2 manipulated category (animate/inanimate) and MSS (1,2,4,8) within subjects, resulting in 8 blocks per participant. Block order was randomized. Each block again consisted of three phases (Figure 3).

**Figure 3.**
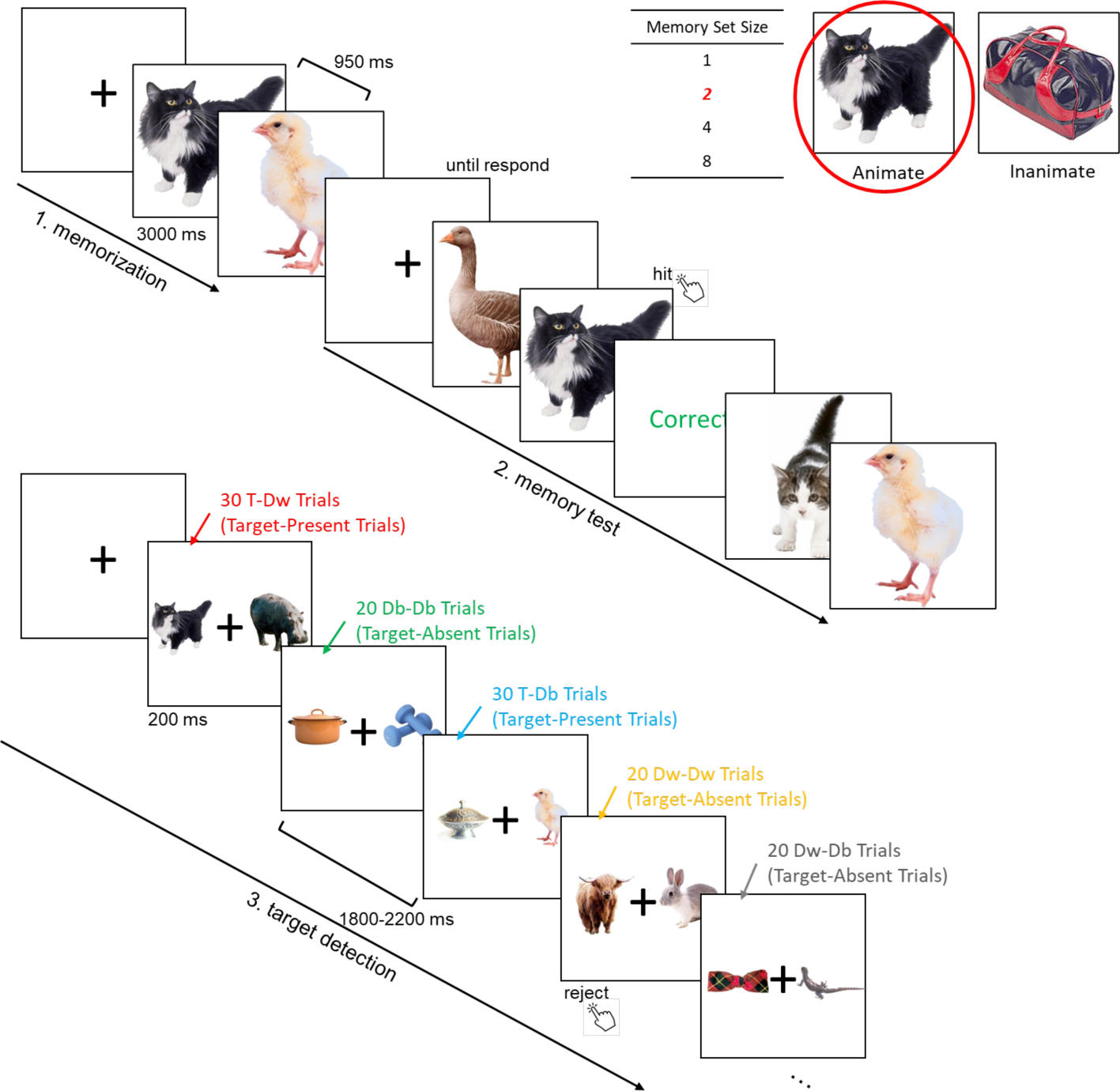
Illustration of the procedure of Experiment 2. The example presents an animate block with two targets (i.e., MSS 2). For target-present trials, a target shown with a within-category distractor condition was abbreviated as T-Dw, a target with a between-category distractor as T-Db; for target-absent trials, trials showing one distractor of the target (i.e., within) category together with one distractor of the other (i.e., between) category was abbreviated as Dw-Db, trials where both distractors were within category as Dw-Dw. Finally, trials where both distractors were between category were abbreviated as Db-Db. Our main analyses focused on T-Dw, T-Db, and Dw-Db conditions, as we hypothesized lateralized responses in these conditions.

Unlike Experiment 1b, the third phase now consisted of a memory search task in which participants were presented with a pair of object images. Participants decided, as quickly as possible, whether the image pair included one of the memorized objects. Each block included a total of 120 trials, presented in random order. In target-present trials (50%), the target was presented together with a within- or a between-category distractor (Figure 3). In target-absent trials (50%), three combinations of two distractors (two within-category distractors, two between-category distractors, one within- and one between-category distractors) were presented with equal probability (Figure 3). Category (animate/inanimate) and target location (left/right visual field) were counterbalanced for each combination. At the beginning of each trial, a black fixation cross appeared for 500 msec, followed by one of the five combinations randomly presented for 200 msec, with a jittered inter-trial interval between 1800 and 2200 msec, during which participants needed to press up arrow key or down arrow key to indicate whether the image pair contained a target or not. Participants were instructed to blink only after making a response.

##### EEG Acquisition and Pre-Processing

The EEG acquisition system and the pre-processing were exactly the same as Experiment 1b. However, trials with correct responses were segmented from -200 to 500 msec relative to the onset of the image pair. Eye movements and other artifacts were removed based on visual inspection. ICA correction was not applied due to the relationship between saccades and visuospatial attention (Kowler et al., 1995). After that, the clean data were re-referenced to the average of all channels. Baseline correction was from -200 to the stimulus onset. All the pre-processing above was completed in Python 3.11 using the MNE toolbox (Gramfort, 2013).

### Statistical Analyses

#### Experiment 1a

All analyses focused on reaction time (RT) to target-absent trials (80% of trials) in the memory search task, for which each image was presented only once. Only correct responses and RTs above 200 msec were included in further analyses. Furthermore, for each participant, RTs beyond 3 standard deviations (SD) from the condition mean were excluded. Under this criterion, a total of 0.260% data points were excluded from further analyses. RT was analyzed in a three-way mixed-design analysis of variance (ANOVA) with one between-(animate-/inanimate-category group) and two within-subjects factors (within-/between-category condition; MSS 1/2/4/8/16). The Greenhouse-Geisser correction was applied to adjust for lack of sphericity (Jennings & Wood, 1976), and only corrected degrees of freedom and *p*-values are reported. Because the three-way interaction was not significant, *F*_(4, 152)_ = 1.228, *p* = 0.277, *η_P_*^2^ = 0.033, we collapsed the data across animate and inanimate groups in all subsequent analyses. Accuracy was >92% in all conditions. The result pattern of accuracy across conditions were in line with the RT results (data not shown).

#### Experiment 1b

The exclusion criteria of behavioral data were identical to those of Experiment 1a. In total, 1.139% of data points were removed. Only correct responses in target-absent trials were included for both behavioral and EEG analysis. Accuracy was >98% in all conditions and the pattern of accuracy across conditions supported the RT results (data not shown).

##### ERP Analyses

The ERP analysis focused on P1, N1, and P2 components over posterior electrodes. We first visually inspected the ERP waveform for each participant. All participants showed ERP waveforms with recognizable P1, N1, and P2 peaks. The time window for each component was defined based on the peak range among the participants (Robinson et al., 2015). P1 peak was observed between 100 to 160 msec after stimulus onset. N1 and P2 peaks were from 160 to 200 msec and from 200 to 300 msec, respectively. Finally, twelve electrodes, P5/P6, P7/P8, PO3/PO4, PO7/PO8, PO9/PO10, and O1/O2, were selected for further analysis as these electrodes showed similar visually evoked activity.

Two-way repeated-measures ANOVA (MSS 1/2/4/8 × within-/between-category conditions) was employed to test for differences in the mean amplitudes of the twelve posterior electrodes across MSS and category conditions, separately for the P1, N1, and P2 components. The Greenhouse-Geisser correction was applied for adjusting for lack of sphericity (Jennings & Wood, 1976), and only corrected degrees of freedom and *p*-value are reported. Then, cluster-based nonparametric permutation tests (Maris & Oostenveld, 2007) were employed to further examine the time courses of the main effects and interaction with 1-ms resolution from 100-300 msec; the range that included the three ERP components.

##### Decoding analyses

Based on the pattern across the twelve electrodes, a linear support vector machine (SVM) was employed to conduct cross-task decoding analysis. The visual oddball task data was used to train an animacy decoder, which was used to decode the object categories in the memory search task between 0 and 600 msec, separately for each participant. Temporal resolution was down-sampled to 100 Hz. The area under the receiver-operator-characteristic curve (AUC) was employed to evaluate the performance on classification, which referred to the probability to distinguish positive and negative classes. As classification metric, it is independent from the classifier threshold, and more robust for imbalanced classes than classification accuracy (Treder, 2020).

#### Experiment 2

The exclusion criteria for removing trials based on behavioral responses were identical to those of Experiment 1. In total, 1.471% of data points were excluded from further analysis. Accuracy was >91% in all conditions and the pattern of accuracy across conditions supported the RT results (data not shown).

##### ERP Analyses

ERP analyses focused on the amplitude of the N2pc component, which was defined by the time window of 200 to 299 msec (Luck & Hillyard, 1994b; Yeh & Peelen, 2022) at two electrode sites PO7/8 (Burra & Kerzel, 2013; Eimer & Kiss, 2008; Kiss et al., 2008; Mazza et al., 2007; Stoletniy et al., 2022). For target-present trials, differences in the N2pc (contra-ipsilateral responses) were tested in a two-way repeated-measures ANOVA (MSS 1/2/4/8 × T-Dw/T-Db category). Finally, cluster-based nonparametric permutation test (Maris & Oostenveld, 2007) was adopted to test the time courses of the main effects and interaction with 1-ms resolution from 100-400 msec.

## Results

### Experiment 1a

Experiment 1a was a behavioral study aimed at replicating previous findings of category effects in memory search (e.g., Drew & Wolfe, 2014) but now using a paradigm that would be suitable to use with EEG (Experiment 1b). To avoid differential repetition effects across conditions (Nosofsky, Cao, et al., 2014), we measured behavioral responses to individually presented distractor objects, with distractor objects making up 80% of trials. Each distractor image was only shown once. We asked: 1) whether search efficiency was modulated by the categorical similarity between the distractors and the objects in the memory set, and 2) whether reaction times for distractors under these conditions would follow the typical log-linear relationship with set size (e.g., Drew & Wolfe, 2014; Wolfe, 2012).

#### Set Size and Category Effects

A two-way repeated-measures ANOVA with RT as dependent variable and memory set size (MSS; 1/2/4/8/16) and category (within-/between-category conditions) as independent variables revealed significant main effects of MSS, *F*_(3.41, 133)_ = 35.521, *p* < 0.001, *η_P_*^2^ = 0.477, and category, *F*_(1, 39)_ = 320.039, *p* < 0.001, *η_P_*^2^ = 0.891. Furthermore, the interaction between MSS and category was significant, *F*_(4, 156)_ = 24.230, *p* < 0.001, *η_P_*^2^ = 0.383. As can be observed in Figure 4A, MSS had a stronger effect (i.e., memory search was less efficient) for within-category than between-category distractors. The simple main effect of set size was significant for both within-, *F*_(4, 156)_ = 54.442, *p* < 0.001, *η_P_*^2^ = 0.583, and between-category conditions, *F*_(4, 156)_ = 10.873, *p* < 0.001, *η_P_*^2^ = 0.218.

**Figure 4.**
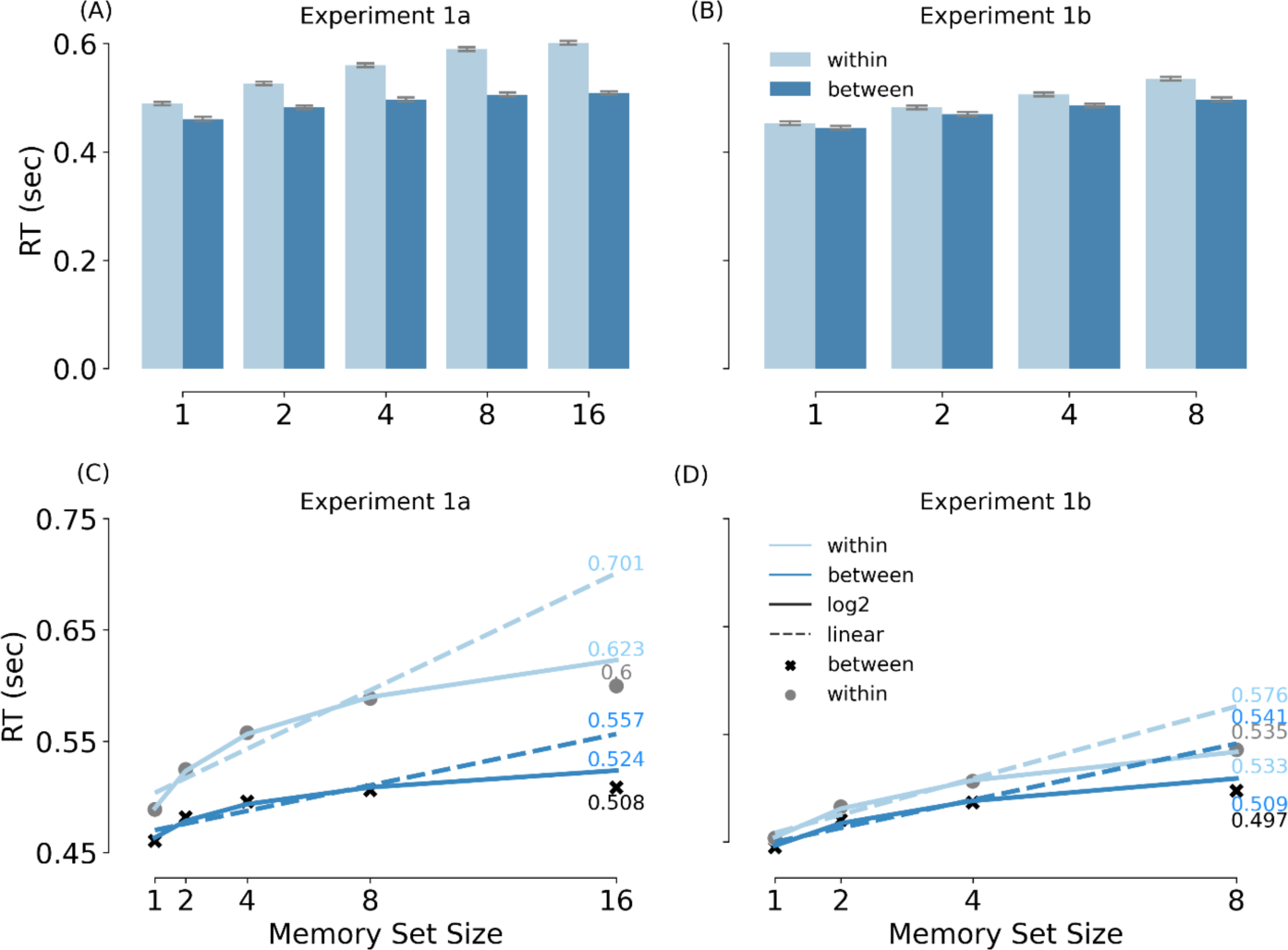
Mean RT (in sec) and fitted models in Experiment 1a and b. (A) and (B) show the mean RTs in Experiment 1a and b, respectively. (C) and (D) show the fitted models in Experiment 1a and b, respectively. The light grey dots and dark grey crosses refer to the observed data in within-category and between-category conditions. The dashed lines represent linear models, while the solid lines represent log-linear models. For all the figures, the error bars represent the standard error of the mean; the light blue represents within-category conditions while the dark blue represents between-category conditions.

#### Linear vs Loglinear models

The increase with MSS visibly displayed a non-linear increase, in line with previous work (Drew & Wolfe, 2014; Wolfe, 2012). To confirm these results statistically, the RTs from set size 1 to 8 were used to predict the performance on set size 16 (Figure 4C), following previous studies (e.g., Drew & Wolfe, 2014). For both category conditions, no significant difference was observed between the observed data and the predicted data based on the log2 model, *t*_(78)_ = 1.406, *p* = 0.164, *d* = 0.314 in within-category condition and *t*_(78)_ = 0.898, *p* = 0.372, *d* = 0.201 in between-category condition. By contrast, the predicted data based on a linear model was significantly higher than the observed data in both within-category condition, *t*_(78)_ = 4.997, *p* < 0.001, *d* = 1.117, and between-category condition, *t*_(78)_ = 2.434, *p* = 0.018, *d* = 0.544.

Finally, we fitted the log-linear model to the observed data using all five set sizes. Confirming the category × MSS interaction observed in the ANOVA, the log-linear slope coefficients for the two category conditions differed significantly, *t*_(39)_ = 9.604, *p* < 0.001, *d* = 1.519, with a steeper slope for within-than between-category distractors.

#### Summary

In this behavioral experiment, we replicated previous findings of a log-linear increase of RT with MSS (e.g., Drew & Wolfe, 2014; Wolfe, 2012). Interestingly, this was observed for distractor objects, which made up 80% of trials. Furthermore, each of these objects was presented only once, excluding the possibility that the set size effect reflected the influence of differential repetition (e.g., items repeating more often in low than high set size conditions; Nosofsky, Cao, et al., 2014; Nosofsky, Cox, et al., 2014). Most importantly for the present purpose, we found a strong category effect on search efficiency: memory search was much more efficient for distractors that were categorically dissimilar to the items in the memory set than for distractors that were of the same category as the items in the memory set (Cunningham & Wolfe, 2012, 2014; Drew & Wolfe, 2014).

##### Experiment 1b

Experiment 1b adopted EEG to test when categorical similarity modulates the processing of the distractor objects. We reasoned that if the between-category advantage is driven by the (proactive) use of categorical attentional templates, this would be observed as a modulation of relatively early visual processing (150-250 ms). By contrast, if the between-category advantage is due to a more efficient search in memory (post visual processing), no such early modulation would be observed. Accordingly, we focused our analysis on two visually evoked event-related potential (ERP) components that emerge within the first 200 ms after stimulus onset: P1 and N1. While the P1 is only modulated by spatial attention, the N1 is modulated by feature-based attention (Hopf et al., 2004; Motter, 1994; but see Zhang & Luck, 2009). Similar to feature-based attention, category-based attention was shown to modulate processing from 150-200 ms after stimulus onset (VanRullen & Thorpe, 2001), with better decoding of attended than unattended categories at this latency (Kaiser et al., 2016). Based on these findings, we expected that a category-based attention mechanism during memory search would similarly modulate the N1 component and increase the accuracy of object category decoding at that latency. Finally, the P2 component was also included in our analyses, based on previous studies implicating the P2 in matching perceptual inputs to memory templates (Dunn et al., 1998; Freunberger et al., 2007; Lefebvre et al., 2005; Luck & Hillyard, 1994a).

#### Behavioral Results

Figure 4B shows the behavioral results of Experiment 1b. These results replicated the findings of Experiment 1a. There were significant main effects of MSS, *F*_(3, 93)_ = 31.591, *p* < 0.001, *η_P_*^2^ = 0.505, and category, *F*_(1, 31)_ = 91.205, *p* < 0.001, *η_P_^2^* = 0.746. As in Experiment 1a, the interaction between MSS and category was significant, *F*_(2.34, 72.61)_ = 6.798, *p* = 0.003, *η_P_^2^* = 0.180. Simple effects of MSS were significant in both within-category condition, *F*_(3, 93)_ = 42.176, *p* < 0.001, *η_P_^2^* = 0.576, and between-category condition, *F*_(3, 93)_ = 14.41, *p* < 0.001, *η_P_^2^* = 0.317. Pairwise comparisons showed significant category effects for all MSSs (*p* < 0.001), except for set size 1 (*p* = 0.175).

The linear/log 2 prediction based on set sizes 1 to 4 demonstrated that the log-linear model was a better fit for both within-category and between-category conditions (Figure 4D): Within category: log-linear vs observed: *t*_(62)_ = -0.121, *p* = 0.904, *d* = 0.030, while linear vs observed: *t*_(62)_ = 2.116, *p* = 0.019, *d* = 0.529 (1-tailed; greater); between category: log-linear vs observed: *t*_(62)_ = 0.575, *p* = 0.568, *d* = 0.144 while linear vs observed: *t*_(62)_ = 1.798, *p* = 0.039, *d* = 0.449 (1-tailed; greater). The slope coefficients between these two conditions (fitting the model on all set sizes) were also significantly different, *t*_(31)_ = 3.488, *p* = 0.001, *d* = 0.617.

#### ERP Results

Separate ANOVAs were run for the three components of interest (P1, N1, P2). There were no significant effects for the P1, MSS effect: *F*_(3, 93)_ = 0.629, *p* = 0.598, *η_P_^2^* = 0.020, category effect: *F*_(1, 31)_ = 1.593, *p* = 0.216, *η_P_^2^* = 0.049, and interaction: *F*_(3, 93)_ = 0.034, *p* = 0.992, *η_P_^2^* = 0.001 (Figure 5B). Importantly, confirming our hypothesis, the N1 showed a significant main effect of category, *F*_(1, 31)_ = 9.185, *p* = 0.005, *η_P_^2^* = 0.229 (Figure 5C). The main effect of MSS, *F*_(3, 93)_ = 2.272, *p* = 0.085, *η_P_^2^* = 0.068, and the interaction between category and MSS, *F*_(3, 93)_ = 0.285, *p* = 0.836, *η_P_^2^* = 0.009, were not significant. Finally, the P2 showed main effects of MSS, *F*_(3, 93)_ = 9.233, *p* < 0.001, *η_P_^2^* = 0.229, and category, *F*_(1, 31)_ = 18.195, *p* < 0.001, *η_P_^2^* = 0.370 (Figure 5D). The interaction between category and MSS was not significant, *F*_(3, 93)_ = 0.650, *p* = 0.585, *η_P_^2^* = 0.021.

**Figure 5.**
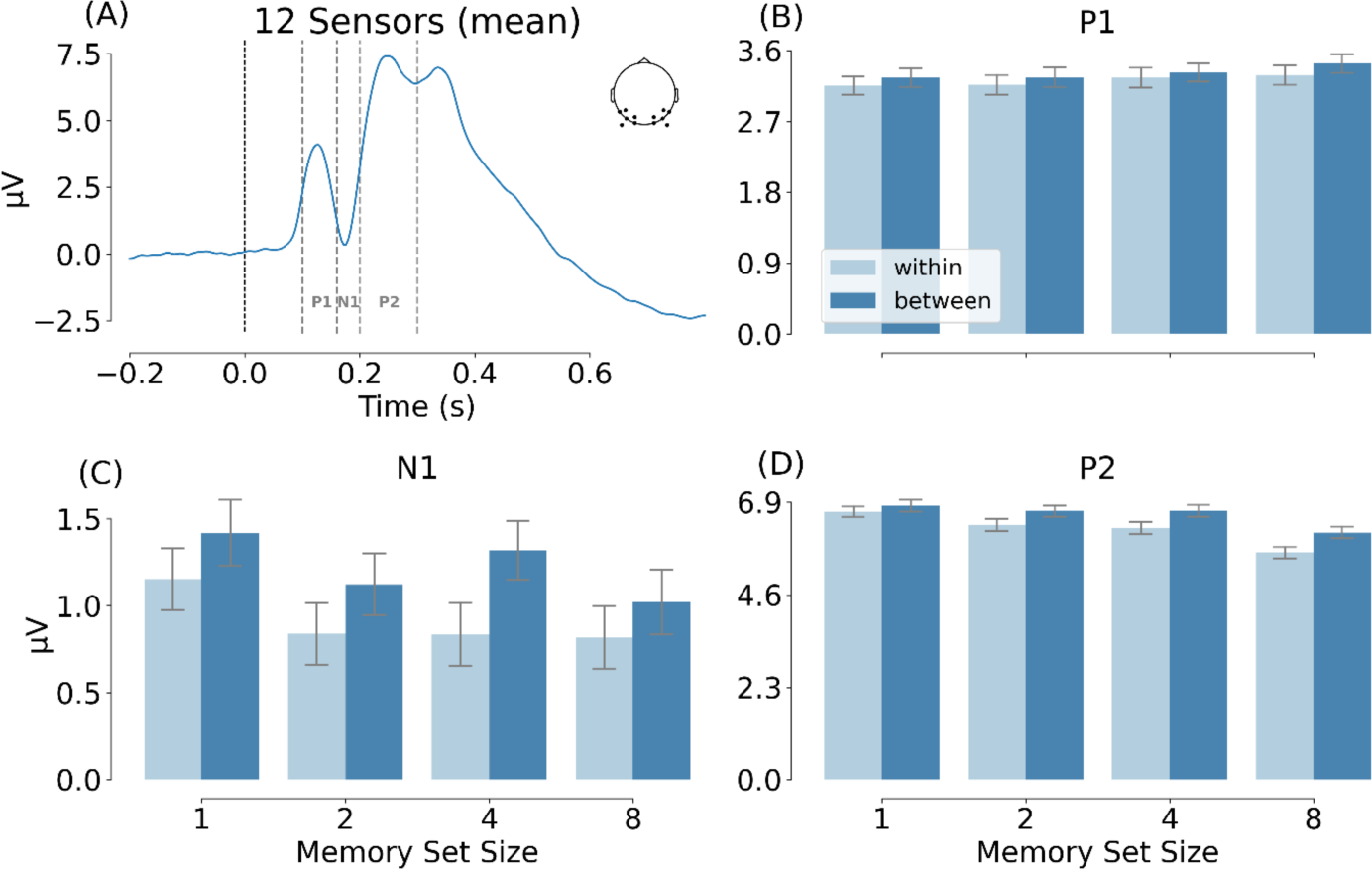
EEG results of Experiment 1b. (A) The mean amplitude based on 12 electrodes (P5/P6, P7/P8, PO3/PO4, PO7/PO8, PO9/PO10, and O1/O2) averaged across all conditions, illustrating the three ERP components of interest. (B), (C), and (D) compare the mean amplitude of the 12 electrodes in the two category and four MSS conditions.

The ERP results were confirmed by a cluster permutation test (Figure 6), showing significant category effects from 152 to 260 msec (cluster-based *p* = 0.001) and significant MSS effects from 188 to 292 msec (cluster-based *p* = 0.002). No interaction effects were found in this analysis.

**Figure 6.**
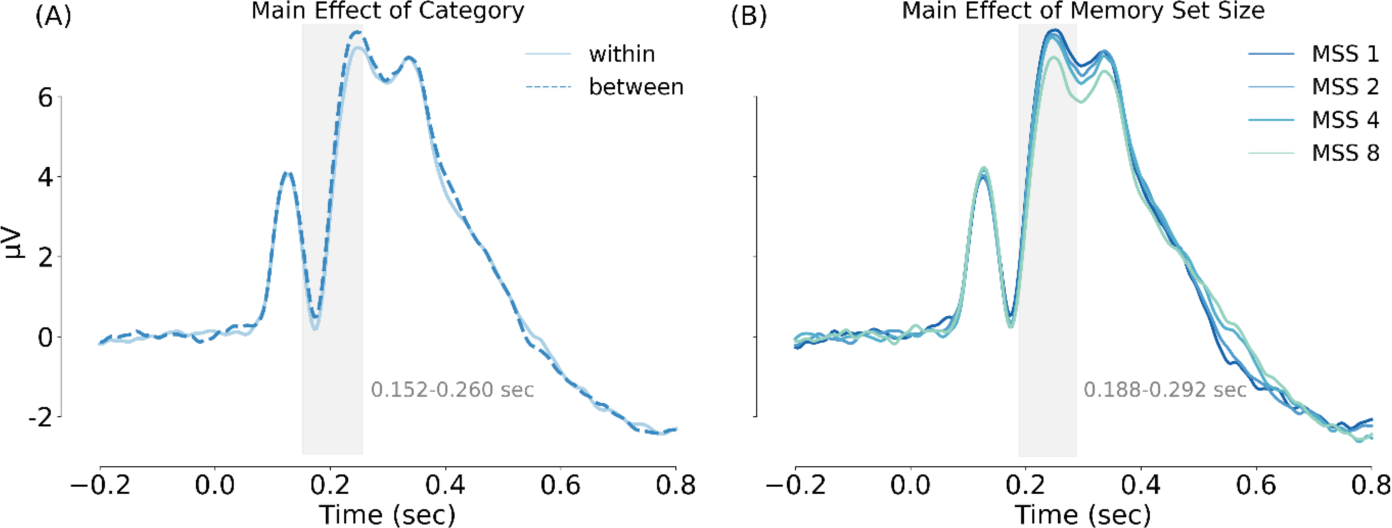
Cluster-based analyses of Experiment 1b. (A) Main effect of category. The light blue solid line represents within-category condition while the dark blue dashed line represents between-category condition. (B) Main effect of MSS. The blue color from dark to light refers to set size 1, 2, 4 and 8, respectively. The grey bars indicate the time windows of significant main effects based on a cluster permutation test.

#### Decoding Results

To test whether attention modulated information about object category, we decoded the category of the distractor objects using a classifier trained on data from a separate experiment that did not involve memory search (see Materials and Methods), following the cross-decoding approach of a previous attention study (Kaiser et al., 2016). Decoding accuracy reflects the representational strength of the distractor objects, rather than the amplitude of the evoked responses. Results showed that AUC scores in all eight conditions reached significance (cluster-based *p* < 0.05) from around 130-150 to 400-580 msec (Figure 7A, 7B), with the first peak at around 170 msec, in line with previous decoding studies (Carlson et al., 2013; Cichy et al., 2014; Kaiser et al., 2016).

**Figure 7.**
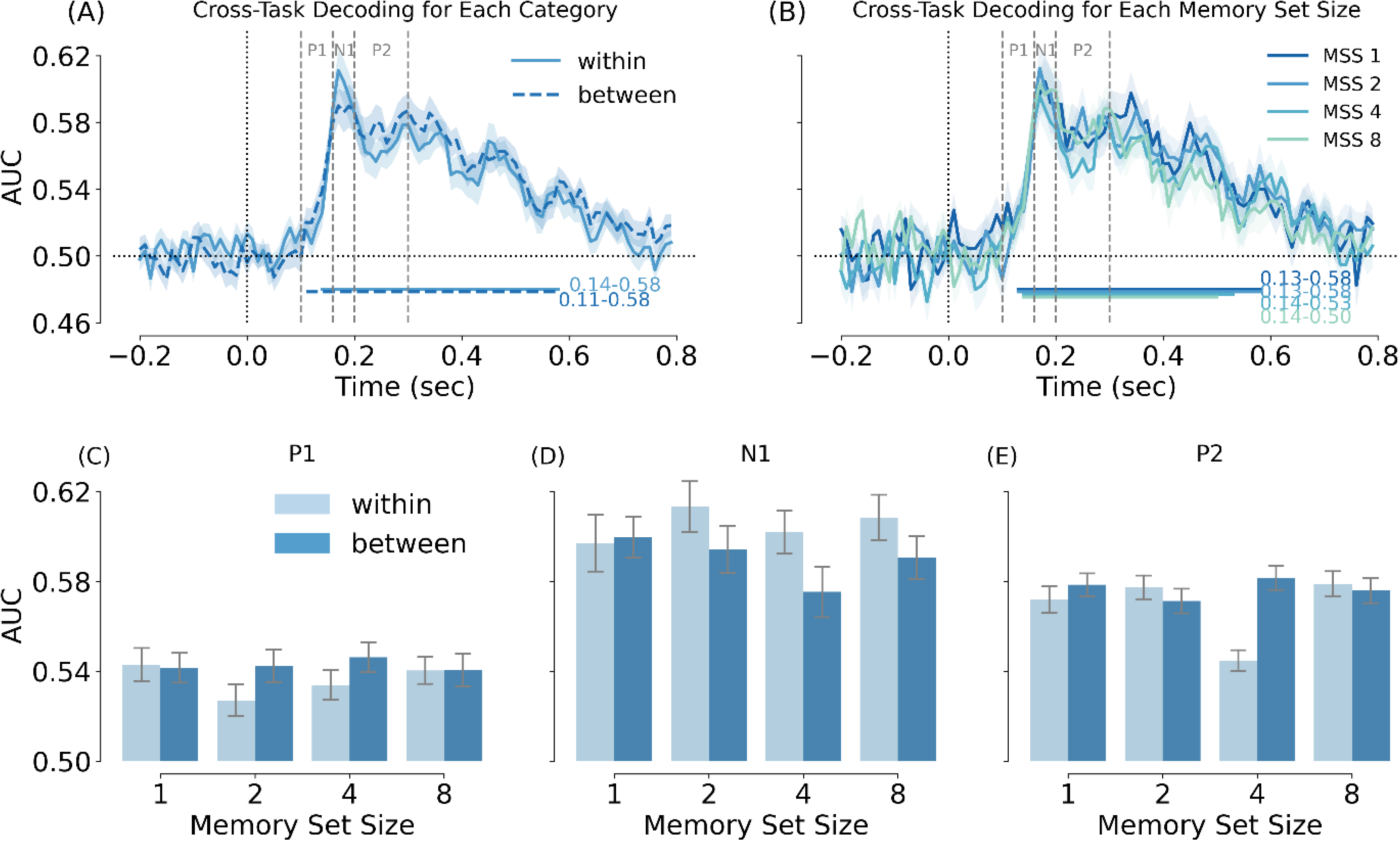
Cross-task decoding in Experiment 1b. (A) AUC as a function of category condition. The light blue solid line represents within-category condition while the dark blue dashed line represents between-category condition. (B) AUC as a function of MSS. The blue color from dark to light refers to set size 1, 2, 4 and 8, respectively. (C), (D), and (E) compare the mean AUC in the two categories and four MSS conditions, averaged across the time points of each component (P1, N1, P2). The main effect of category was significant for the N1 time window (panel D).

Next, we averaged decoding accuracy across the time window of each ERP component and tested these using repeated-measures ANOVAs. In line with the ERP results, no significant effects were observed for the P1 time window (*p* > 0.335, for all tests; Figure 7C). Interestingly, the N1 showed a significant main effect of category, *F*_(1, 31)_ = 4.516, *p* = 0.042, *η_P_^2^* = 0.127, with better decoding for within-than between-category distractors (Figure 7D). The main effect of MSS, and the interaction between MSS and category, were not significant (*p* > 0.678, for all tests). Finally, no significant effects were observed for the P2 time window (*p* > 0.109, for all tests; Figure 7E).

#### Summary

The behavioral results of Experiment 1b replicated those of Experiment 1a, again showing that memory search was more efficient for between-than within-category distractors. The ERP results showed that category membership modulated the visually evoked N1 component (160-200 ms) as well as the subsequent P2 component (200-300 ms), while set size only modulated the P2 component. The cluster permutation test confirmed these results, showing a relatively early category effect, from 152 to 260 msec, while set size effects emerged from 188 to 292 msec. Finally, the decoding results of the N1 window showed that object category decoding was higher for within-category distractors than between-category distractors. Altogether, these results provide evidence for category-level attentional modulation during a memory search task. Distractor objects matching the category of the memory set received more processing than non-matching distractor objects, demonstrated both by a differential evoked responses (VanRullen & Thorpe, 2001) and more accurate categorical representation (Kaiser et al., 2016) around 160-200 ms after onset.

##### Experiment 2

In Experiment 2, we followed up on the findings of Experiment 1b, testing whether category-matching distractors attract spatial attention. By having participants search for targets in two-object displays (Figure 3), we could measure the allocation of spatial attention using the lateralized N2pc EEG component: previous studies showed that template-matching objects (e.g., targets) during visual search attract spatial attention, eliciting an N2pc (Eimer, 1996; Luck et al., 2000). This target-elicited N2pc is reduced when a target appears together with a distractor that partially matches the template (Nako et al., 2016; Wu et al., 2016; Yeh et al., 2019; Yeh & Peelen, 2022). Therefore, if participants adopted a category-level attentional template in our memory search task, we expected the N2pc to be reduced when a target appeared next to a within-category (“T-Dw”) as compared to a between-category (“T-Db”) distractor. For the same reason, we expected to observe an N2pc in response to a within-category distractor (“Dw”) when shown together with a between-category (“Db”) distractor.

#### Behavioral Results

Figure 8A shows the RT results for target-present trials. A two-way repeated-measures ANOVA (MSS; 1/2/4/8 × category; T-Dw/T-Db) showed a main effect of MSS, *F*_(2.03, 62.88)_ = 104.097, *p* < 0.001, *η_P_^2^* = 0.771, a main effect of category, *F*_(1, 31)_ = 32.212, *p* < 0.001, *η_P_^2^* = 0.510, and an interaction between MSS and category, *F*_(3, 93)_ = 4.568, *p* < 0.001, *η_P_^2^* = 0.128. Simple main effects of MSS were also significant within both the T-Dw condition, *F*_(2.23, 69.1)_ = 88.3, *p* < 0.001, *η_P_^2^* = 0.576, and the T-Db condition, *F*_(1.93, 59.9)_ = 92.5, *p* < 0.001, *η_P_^2^* = 0.749. Simple main effects of category were observed in MSS 4 and 8, *F*_(1, 31)_ = 20.4, *p* < 0.001, *η_P_^2^* = 0.397 and *F*_(1, 31)_ = 11.5, *p* = 0.002, *η_P_^2^* = 0.271, but not in MSS 1 and 2, *F*_(1, 31)_ = 0.65, *p* = 0.426, *η_P_^2^* = 0.021 and *F*_(1, 31)_ = 1.55, *p* = 0.222, *η_P_^2^* = 0.048.

For target-absent trials (Figure 8B), a two-way repeated-measures ANOVA (MSS 1/2/4/8 × Dw-Db/Dw-Dw/Db-Db category) showed a main effect of MSS, *F*_(3,93)_ = 99.905, *p* < 0.001, *η_P_^2^* = 0.763, a main effect of category, *F*_(1.61,50.05)_ = 88.455, *p* < 0.001, *η_P_^2^* = 0.740, and an interaction between set size and category, *F*_(3.7,114.82)_ = 12.108, *p* < 0.001, *η_P_^2^* = 0.281. Significant main effects of MSS were observed in all three category conditions (Dw-Db: *F*_(3, 93)_ = 95.57, *p* < 0.001, *η_P_^2^* = 0.755; Dw-Dw: *F*_(3, 93)_ = 75.906, *p* < 0.001, *η_P_^2^* = 0.710; Db-Db: *F*_(3, 93)_ = 34.368, *p* < 0.001, *η_P_^2^* = 0.526).

**Figure 8.**
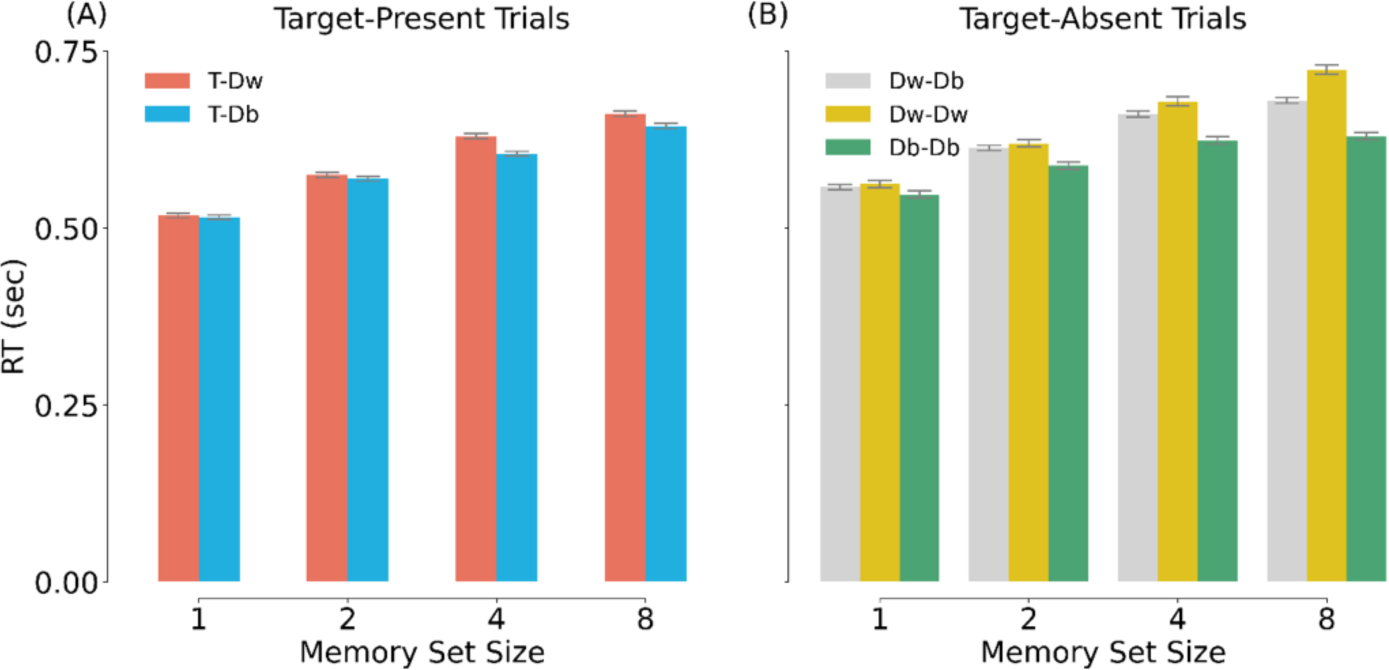
Mean RT (in sec) in Experiment 2. (A) RTs in target-present trials. The red bars represent the within-category condition (T-Dw; target shown next to a within-category distractor) while the blue bars represent the between-category condition (T-Db; target shown next to a between-category distractor). (B) RTs in target-absent trials. The grey, yellow, and green bars represent Dw-Db, Dw-Dw, and Db-Db trials.

#### ERP Results

*N2pc induced by targets.* In a first analysis, we wanted to verify that the targets in our experiment evoked a reliable N2pc. Averaged across conditions, we observed a strong N2pc, with a more negative response contralateral vs ipsilateral to the target from around 200 ms after stimulus onset (Figure 9A). Next, we averaged the amplitude of the evoked response in the N2pc time window (200-299 ms after onset) and tested the N2pc effect for each set size (Figure 9B). This analysis revealed a significant N2pc for each set size (*p* < 0.005, for all tests).

**Figure 9.**
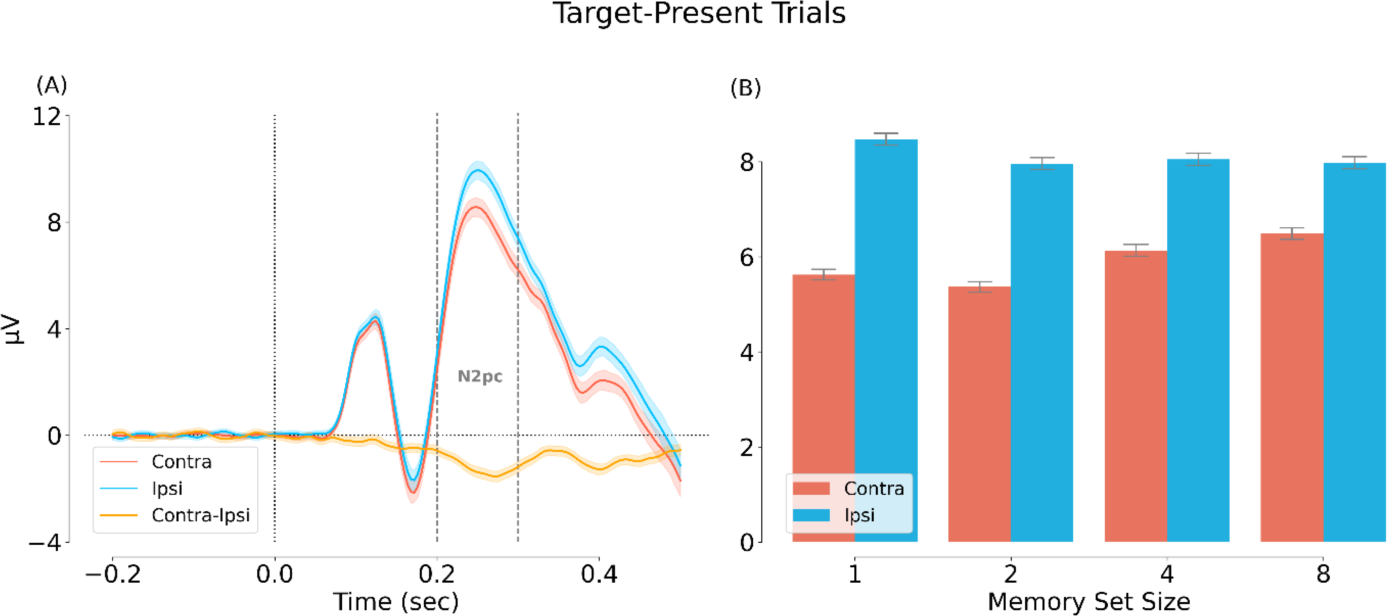
Main effect of visual field location, showing a target related N2pc. (A) Main effect of contra/ipsi-lateral visual field. (B) Comparison of the contra/ipsi-lateral amplitude averaged across the N2pc time window (200-299 msec).

Having established a reliable target-related N2pc, we then tested how this effect was modulated by MSS and category through a two-way repeated-measures ANOVA with the N2pc as dependent variable (contra-ipsi, averaged across 200-299 ms) and MSS (1/2/4/8) and category (T-Dw/T-Db) as independent variables. Results are shown in Figure 10A. The main effects of MSS and category were significant, MSS: *F*_(3, 93)_ = 8.672, *p* < 0.001, *η_P_^2^* = 0.219, category: *F*_(1, 31)_ = 49.580, *p* < 0.001, *η_P_^2^* = 0.615. The interaction between MSS and category was also significant, *F*_(3, 93)_ = 4.822, *p* = 0.004, *η_P_^2^* = 0.135. Following up on the interaction, we found that the simple main effect of MSS was significant in the T-Dw condition, *F*_(3, 93)_ = 11.4, *p* < 0.001, *η_P_^2^* = 0.268, but not in the T-Db condition, *F*_(3, 93)_ = 1.87, *p* = 0.14, *η_P_^2^* = 0.057. Furthermore, the effect of category was significant for MSS 4 and 8, *F*_(1, 31)_ = 20.1, *p* < 0.001, *η_P_^2^* = 0.394 and *F*_(1, 31)_ = 19.1, *p* < 0.001, *η_P_^2^* = 0.381, but not for MSS 1 and 2, *F*_(1, 31)_ = 0.64, *p* = 0.43, *η_P_^2^* = 0.02 and *F*_(1, 31)_ = 1.81, *p* = 0.188, *η_P_^2^* = 0.055.

**Figure 10.**
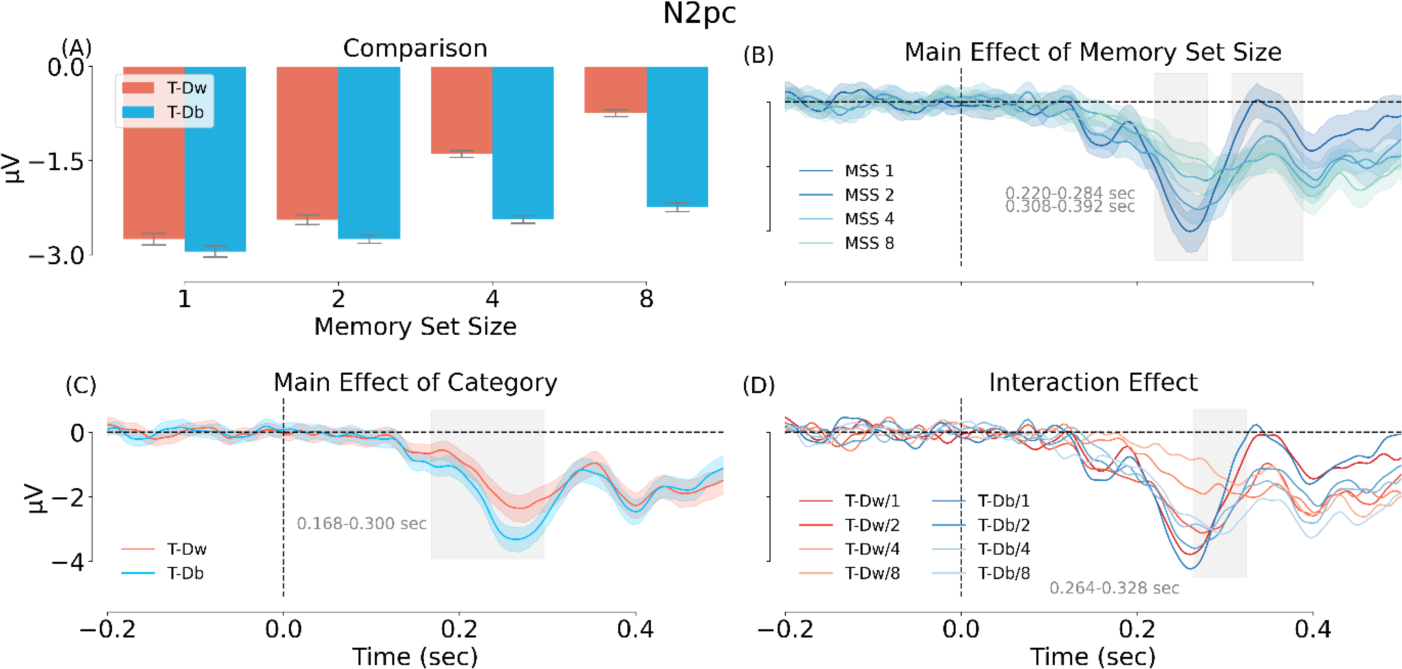
ERP results of Experiment 2. (A) Comparison of the mean N2pc amplitude in the two categories and four MSS conditions. (B) Main effect of MSS in a cluster analysis. (C) Main effect of category in a cluster analysis. (D) Interaction of MSS and category in a cluster analysis. Different colors from dark to light refer to MSS 1 to 8. T-Dw = a target shown next to a within-category distractor, T-Db = a target shown next to a between-category distractor.

These results were confirmed by cluster permutation tests (Figures 10B and 10C), which showed significant MSS effects from 220 to 284 msec and 308 to 392 msec (both cluster-based *p* = 0.001) and significant category effects from 168 to 300 msec (cluster-based *p* = 0.001). The interaction of the two effects was significant from 264 to 328 msec (cluster-based *p* = 0.013).

Together, these results confirm our first hypothesis, that the target-induced N2pc is reduced in the presence of a within-category distractor. Mirroring the behavioral results, this reduction was stronger for larger set size.

*N2pc induced by distractors.* Next, we tested our second hypothesis, that of an N2pc induced by a within-category (“Dw”) versus between-category (“Db”) distractors. Dw-Dw and Db-Db trials were not included in the analysis because there was, by definition, no lateralized attentional bias in these two conditions. Results confirmed our hypothesis: we observed a significant difference between contra- and ipsi-lateral responses from around 200 ms after stimulus onset (Figure 11A). Averaging responses across the N2pc time window (200 and 299 msec) revealed a significant N2pc, *F*_(1, 31)_ = 121.917, *p* < 0.001, *η_P_^2^* = 0.797. The N2pc did not differ significantly across set size, *F*_(3, 93)_ = 1.474, *p* = 0.227, *η_P_^2^* = 0.045 (Figure 11B).

**Figure 11.**
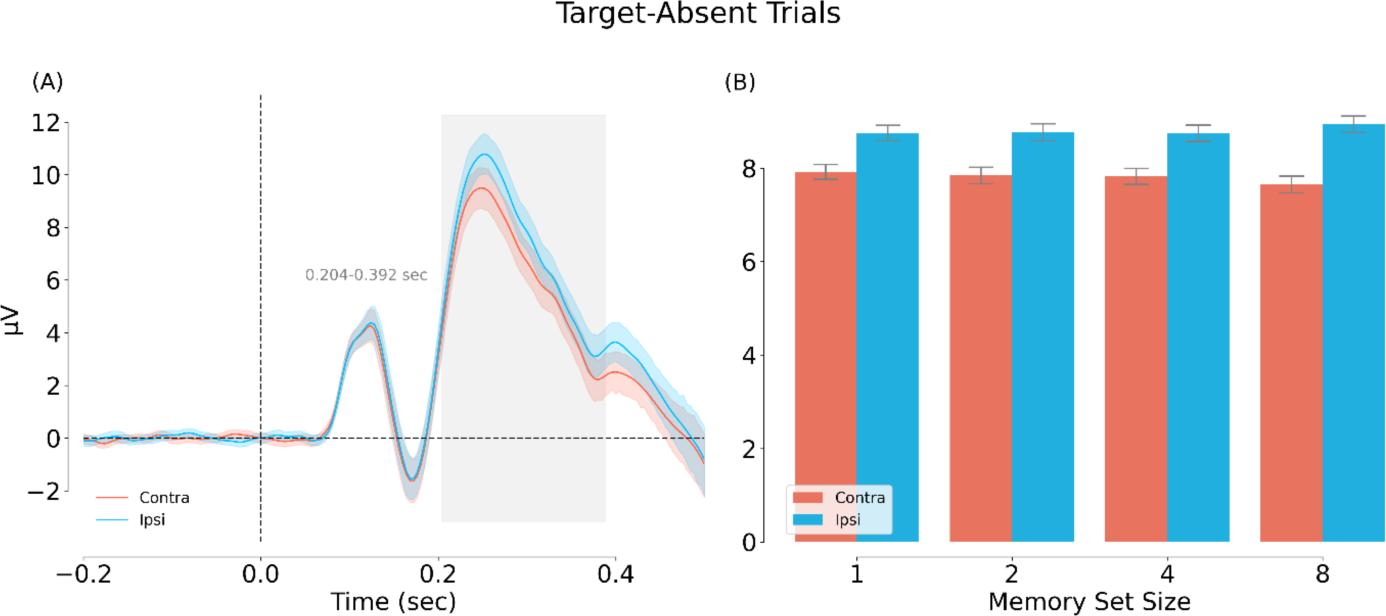
Main effect of visual field location for the Dw-Db condition (target-absent trials). In this analysis, contralateral is relative to the location of the within-category distractor (Dw). (A) Main effect of contra/ipsi-lateral visual field based on a cluster permutation test (cluster-based *p* = 0.001). (B) Comparison of the contra/ipsi-lateral amplitude averaged across the N2pc time window (200-299 msec).

#### Summary

The results of Experiment 2 showed that the target-elicited N2pc was reduced when the target was shown next to a same-category distractor. Furthermore, spatial attention (indexed by the N2pc component) was directed towards distractors that matched the category of the memory set. Altogether, these results provide evidence that participants formed a categorical attentional template, with spatial attention being directed to distractor objects belonging to the category of the memory set.

## Discussion

In three experiments, we investigated the role of attention in memory search. In an online behavioral experiment (Experiment 1a), participants memorized target objects; then, during a memory search phase, they viewed one object at a time and decided whether the object was part of the memorized set of objects. The memory set always consisted of objects from a single category (animate or inanimate objects), while the distractor objects could be from this or another category. Our analyses focused on responses to these distractor objects (80% of trials) as a function of memory set size (1, 2, 4, 8, 16) and category (same or different category as the memory set). Results showed that memory search was much more efficient for between-category than within-category distractors, replicating earlier work (Cunningham & Wolfe, 2012, 2014; Drew & Wolfe, 2014). Using EEG (Experiment 1b), we tested whether this increased efficiency could be explained by attentional modulation at the level of object category. Results confirmed this hypothesis, showing that early visual object processing was modulated by the category membership of the distractor: We found a larger N1 in response to distractors from the target category compared to distractors from a different category.

Furthermore, decoding analyses showed that within-category distractors were more strongly represented than between-category distractors at this latency. The results of Experiment 1 are in line with the attentional modulation of visual processing observed in single-target detection tasks (Kaiser et al., 2016; VanRullen & Thorpe, 2001). This modulation is much earlier than the typical time window of memory retrieval, which usually starts after ∼300 msec (Curran & Hancock, 2007; Noh et al., 2018; Rugg et al., 1998; Rugg & Curran, 2007). In Experiment 2, we presented two objects simultaneously to test whether spatial attention (indexed by the N2pc component) was guided to the location of template-matching objects (Battistoni et al., 2020; Eimer, 2014; Hopf et al., 2004; Wolfe, 2021). We found that spatial attention was directed towards distractor objects that were of the same category as the items in the memory set. Together, the results provide evidence that participants spontaneously use the shared category of the memory items to form a category-level attentional template. By allocating more attentional resources to the features (N1) and location (N2pc) of the target category, they were able to efficiently reject between-category distractors before commencing search in long-term memory.

The behavioral results of all experiments (Figure 4; Figure 8) and the target related N2pc results of Experiment 2 (Figure 10A) revealed that category interacted with memory set size, such that the categorical modulation became stronger with increasing set size. This result can be explained in at least two ways. First, it is possible that participants only started to use an attentional template-based strategy when memory search became effortful (i.e., with high set size). Alternatively, participants may have adopted an attentional template-based strategy for all set sizes, but the specificity of the template varied with the memory set size. For a set size of two, the template may have been specific to the subcategories of the two targets (e.g., a cat and a horse). In that case, distractor objects from other subcategories (e.g., reptile, bird) may not have provided a strong match to the template. Instead, for higher set sizes, a larger number of subcategories made up the memory set. In that case, the features that the subcategories had in common were more likely to generalize to the distractor objects from the same superordinate category (e.g., animals). Future studies could systematically manipulate the similarity of the subcategories within the memory set to distinguish between these accounts.

The interaction between set size and category also shows that set size more strongly affected the rejection of within-category than between-category distractors. This suggests that within-category distractors activated a memory search process, while between-category distractors did so only weakly. Of note, the effect of set size was still significant for between-category distractors in all behavioral analyses, suggesting that categorical attention was not fully preventing between-category distractors from entering the memory search process.

Interestingly, not all analyses showed an interaction between set size and category. Specifically, the modulation of visual distractor processing in Experiment 1b (Figure 5C; 7D) and the distractor evoked N2pc in Experiment 2 (Figure 11B) only showed a main effect of category. It is possible that the absence of an interaction in these analyses reflected a lack of power (e.g., see the weak trend towards an interaction in Figure 11B). Alternatively, the interaction may reflect a true dissociation between these measures. For example, in Experiment 2, if we assume that the within-category distractor only weakly matched the template when set size was low (e.g., because the template was specific to the subcategories in the memory set) these distractors would not have provided strong competition when shown next to a target, resulting in the category x set size interaction observed for the target-related N2pc. However, when the same within-category distractor is shown next to a between-category distractor, it would still provide a relatively better match to the template than the between-category distractor, and thus attract spatial attention even in the low set size conditions.

Our findings raise the question of what features the categorical template consists of, and whether categorical templates are specific to the categories used here. Animate and inanimate objects differ in terms of mid-, and high-level visual features (e.g., Jozwik et al., 2022; Long et al., 2018; Proklova et al., 2016; Thorat et al., 2019) and it has been proposed that the human visual system is particularly sensitive to these category-diagnostic features (New et al., 2007), as also reflected in the animate-inanimate organization of the ventral visual cortex (Chao et al., 1999; Grill-Spector & Weiner, 2014; Kriegeskorte et al., 2008; Thorat et al., 2019). This raises the possibility that categorical attention in memory search, as revealed here, is specific to the distinction between animate and inanimate objects. Future studies will need to test whether our results generalize to other categorical distinctions (e.g., fruit vs non-fruit). We anticipate that results are most likely to generalize to categories that, like animals, are highly familiar and that are characterized by diagnostic visual features (Battistoni et al., 2017).

To conclude, our study reveals a crucial role of attention in memory search. When observers look for multiple objects at the same time, they can use the objects’ shared categorical features to direct attention at that level, leading to the efficient rejection of distractor objects belonging to other categories (Figure 1B).

## References

Bansal, A. K., Madhavan, R., Agam, Y., Golby, A., Madsen, J. R., & Kreiman, G. (2014). Neural Dynamics Underlying Target Detection in the Human Brain. Journal of Neuroscience, 34(8), 3042–3055. 10.1523/JNEUROSCI.3781-13.2014

Battistoni, E., Kaiser, D., Hickey, C., & Peelen, M. V. (2020). The time course of spatial attention during naturalistic visual search. Cortex, 122, 225–234. 10.1016/j.cortex.2018.11.018

Battistoni, E., Stein, T., & Peelen, M. V. (2017). Preparatory attention in visual cortex: Preparatory attention in visual cortex. Annals of the New York Academy of Sciences, 1396(1), 92–107. 10.1111/nyas.13320

Brady, T. F., Konkle, T., Alvarez, G. A., & Oliva, A. (2008). Visual long-term memory has a massive storage capacity for object details. Proceedings of the National Academy of Sciences of the United States of America, 105(38), 14325–14329. 10.1073/pnas.0803390105

Burra, N., & Kerzel, D. (2013). Attentional capture during visual search is attenuated by target predictability: Evidence from the N2pc, Pd, and topographic segmentation: Saliency and target predictability. Psychophysiology, 50(5), 422–430. 10.1111/psyp.12019

Chao, L. L., Haxby, J. V., & Martin, A. (1999). Attribute-based neural substrates in temporal cortex for perceiving and knowing about objects. Nature neuroscience, 2(10), 913–919.

Chelazzi, L., Miller, E. K., Duncan, J., & Desimone, R. (1993). A neural basis for visual search in inferior temporal cortex. Nature, 363(6427), 345–347. 10.1038/363345a0

Cunningham, C. A., & Wolfe, J. M. (2012). Lions or tigers or bears: Oh my! Hybrid visual and memory search for categorical targets. Visual Cognition, 20(9), 1024–1027. 10.1080/13506285.2012.726455

Cunningham, C. A., & Wolfe, J. M. (2014). The role of object categories in hybrid visual and memory search. Journal of Experimental Psychology: General, 143(4), 1585–1599. 10.1037/a0036313

Curran, T., & Hancock, J. (2007). The FN400 indexes familiarity-based recognition of faces. NeuroImage, 36(2), 464–471. 10.1016/j.neuroimage.2006.12.016

Desimone, R., & Duncan, J. (1995). Neural Mechanisms of Selective Visual Attention. Annual Review of Neuroscience, 18(1), 193–222. 10.1146/annurev.ne.18.030195.001205

Drew, T., & Wolfe, J. M. (2014). Hybrid search in the temporal domain: Evidence for rapid, serial logarithmic search through memory. *Attention, Perception*, & Psychophysics, 76(2), 296–303. 10.3758/s13414-013-0606-y

Dunn, B. R., Dunn, D. A., Languis, M., & Andrews, D. (1998). The Relation of ERP Components to Complex Memory Processing. Brain and Cognition, 36(3), 355–376. 10.1006/brcg.1998.0998

Eimer, M. (1996). The N2pc component as an indicator of attentional selectivity. Electroencephalography and Clinical Neurophysiology, 99(3), 225–234. 10.1016/0013-4694(96)95711-9

Eimer, M., & Kiss, M. (2008). Involuntary Attentional Capture is Determined by Task Set: Evidence from Event-related Brain Potentials. Journal of Cognitive Neuroscience, 20(8), 1423–1433. 10.1162/jocn.2008.20099

Freunberger, R., Klimesch, W., Doppelmayr, M., & Höller, Y. (2007). Visual P2 component is related to theta phase-locking. Neuroscience Letters, 426(3), 181–186. 10.1016/j.neulet.2007.08.062

Gramfort, A. (2013). MEG and EEG data analysis with MNE-Python. Frontiers in Neuroscience, 7. 10.3389/fnins.2013.00267

Grill-Spector, K., & Weiner, K. S. (2014). The functional architecture of the ventral temporal cortex and its role in categorization. Nature Reviews Neuroscience, 15(8), 536–548.

Hopf, J.-M., Boelmans, K., Schoenfeld, M. A., Luck, S. J., & Heinze, H.-J. (2004). Attention to Features Precedes Attention to Locations in Visual Search: Evidence from Electromagnetic Brain Responses in Humans. The Journal of Neuroscience, 24(8), 1822–1832. 10.1523/JNEUROSCI.3564-03.2004

Houtkamp, R., & Roelfsema, P. R. (2006). The effect of items in working memory on the deployment of attention and the eyes during visual search. Journal of Experimental Psychology: Human Perception and Performance, 32(2), 423–442. 10.1037/0096-1523.32.2.423

Jennings, J., & Wood, C. C. (1976). The epsilon-Adjustment Procedure for Repeated-Measures Analyses of Variance. Psychophysiology, 13(3), 277–278. 10.1111/j.1469-8986.1976.tb00116.x

Jozwik, K. M., Najarro, E., Van Den Bosch, J. J., Charest, I., Cichy, R. M., & Kriegeskorte, N. (2022). Disentangling five dimensions of animacy in human brain and behaviour. Communications Biology, 5(1), 1247.

Kaiser, D., Oosterhof, N. N., & Peelen, M. V. (2016). The Neural Dynamics of Attentional Selection in Natural Scenes. Journal of Neuroscience, 36(41), 10522–10528. 10.1523/JNEUROSCI.1385-16.2016

Kiss, M., Van Velzen, J., & Eimer, M. (2008). The N2pc component and its links to attention shifts and spatially selective visual processing. Psychophysiology, 45(2), 240–249. 10.1111/j.1469-8986.2007.00611.x

Kowler, E., Anderson, E., Dosher, B., & Blaser, E. (1995). The role of attention in the programming of saccades. Vision Research, 35(13), 1897–1916. 10.1016/0042-6989(94)00279-U

Kriegeskorte, N., Mur, M., Ruff, D. A., Kiani, R., Bodurka, J., Esteky, H., … & Bandettini, P. A. (2008). Matching categorical object representations in inferior temporal cortex of man and monkey. Neuron, 60(6), 1126–1141.

Lefebvre, C. D., Marchand, Y., Eskes, G. A., & Connolly, J. F. (2005). Assessment of working memory abilities using an event-related brain potential (ERP)-compatible digit span backward task. Clinical Neurophysiology, 116(7), 1665–1680. 10.1016/j.clinph.2005.03.015

Long, B., Yu, C. P., & Konkle, T. (2018). Mid-level visual features underlie the high-level categorical organization of the ventral stream. Proceedings of the National Academy of Sciences, 115(38), E9015–E9024.

Luck, S. J., & Hillyard, S. A. (1994a). Electrophysiological correlates of feature analysis during visual search. Psychophysiology, 31(3), 291–308. 10.1111/j.1469-8986.1994.tb02218.x

Luck, S. J., & Hillyard, S. A. (1994b). Spatial filtering during visual search: Evidence from human electrophysiology. Journal of Experimental Psychology: Human Perception and Performance, 20(5), 1000–1014. 10.1037/0096-1523.20.5.1000

Luck, S. J., Woodman, G. F., & Vogel, E. K. (2000). Event-related potential studies of attention. Trends in Cognitive Sciences, 4(11), 432–440. 10.1016/S1364-6613(00)01545-X

Maris, E., & Oostenveld, R. (2007). Nonparametric statistical testing of EEG- and MEG-data. Journal of Neuroscience Methods, 164(1), 177–190. 10.1016/j.jneumeth.2007.03.024

Mazza, V., Turatto, M., Umiltà, C., & Eimer, M. (2007). Attentional selection and identification of visual objects are reflected by distinct electrophysiological responses. Experimental Brain Research, 181(3), 531–536. 10.1007/s00221-007-1002-4

Motter, B. (1994). Neural correlates of attentive selection for color or luminance in extrastriate area V4. The Journal of Neuroscience, 14(4), 2178–2189. 10.1523/JNEUROSCI.14-04-02178.1994

Nako, R., Grubert, A., & Eimer, M. (2016). Category-based guidance of spatial attention during visual search for feature conjunctions. Journal of Experimental Psychology: Human Perception and Performance, 42(10), 1571–1586. 10.1037/xhp0000244

New, J., Cosmides, L., & Tooby, J. (2007). Category-specific attention for animals reflects ancestral priorities, not expertise. Proceedings of the National Academy of Sciences, 104(42), 16598–16603.

Noh, E., Liao, K., Mollison, M. V., Curran, T., & Sa, V. R. de. (2018). Single-Trial EEG Analysis Predicts Memory Retrieval and Reveals Source-Dependent Differences. Frontiers in Human Neuroscience, 12, 258. 10.3389/fnhum.2018.00258

Nosofsky, R. M., Cao, R., Cox, G. E., & Shiffrin, R. M. (2014). Familiarity and categorization processes in memory search. Cognitive Psychology, 75, 97–129. 10.1016/j.cogpsych.2014.08.003

Nosofsky, R. M., Cox, G. E., Cao, R., & Shiffrin, R. M. (2014). An exemplar-familiarity model predicts short-term and long-term probe recognition across diverse forms of memory search. *Journal of Experimental Psychology: Learning*, Memory, and Cognition, 40(6), 1524–1539. 10.1037/xlm0000015

Olivers, C. N. L., Peters, J., Houtkamp, R., & Roelfsema, P. R. (2011). Different states in visual working memory: When it guides attention and when it does not. In Trends in Cognitive Sciences (Vol. 15). 10.1016/j.tics.2011.05.004

Ort, E., & Olivers, C. N. (2020). The capacity of multiple-target search. Visual Cognition, 28(5-8), 330–355

Peirce, J., Gray, J. R., Simpson, S., MacAskill, M., Höchenberger, R., Sogo, H., Kastman, E., & Lindeløv, J. K. (2019). PsychoPy2: Experiments in behavior made easy. Behavior Research Methods, 51(1), 195–203. 10.3758/s13428-018-01193-y

Proklova, D., Kaiser, D., & Peelen, M. V. (2016). Disentangling representations of object shape and object category in human visual cortex: The animate–inanimate distinction. Journal of cognitive neuroscience, 28(5), 680–692.

Robinson, A. K., Reinhard, J., & Mattingley, J. B. (2015). Olfaction Modulates Early Neural Responses to Matching Visual Objects. Journal of Cognitive Neuroscience, 27(4), 832–841. 10.1162/jocn_a_00732

Rugg, M. D., & Curran, T. (2007). Event-related potentials and recognition memory. Trends in Cognitive Sciences, 11(6), 251–257. 10.1016/j.tics.2007.04.004

Rugg, M. D., Mark, R. E., Walla, P., Schloerscheidt, A. M., Birch, C. S., & Allan, K. (1998). Dissociation of the neural correlates of implicit and explicit memory. Nature, 392(6676), 595–598. 10.1038/33396

Sternberg, S. (1966). High-Speed Scanning in Human Memory. Science, 153(3736), 652–654. 10.1126/science.153.3736.652

Stoletniy, A. S., Alekseeva, D. S., Babenko, V. V., Anokhina, P. V., & Yavna, D. V. (2022). The N2pc Component in Studies of Visual Attention. Neuroscience and Behavioral Physiology, 52(8), 1299–1309. 10.1007/s11055-023-01359-y

Thorat, S., Proklova, D., & Peelen, M. V. (2019). The nature of the animacy organization in human ventral temporal cortex. Elife, 8, e47142.

Treder, M. S. (2020). MVPA-Light: A Classification and Regression Toolbox for Multi-Dimensional Data. Frontiers in Neuroscience, 14, 289. 10.3389/fnins.2020.00289

van Moorselaar, D., Theeuwes, J., & Olivers, C. N. L. (2014). In competition for the attentional template: Can multiple items within visual working memory guide attention? Journal of Experimental Psychology: Human Perception and Performance, 40(4), 1450–1464. 10.1037/a0036229

VanRullen, R., & Thorpe, S. J. (2001). The Time Course of Visual Processing: From Early Perception to Decision-Making. Journal of Cognitive Neuroscience, 13(4), 454–461. 10.1162/08989290152001880

Wolfe, J. M. (2012). Saved by a Log: How Do Humans Perform Hybrid Visual and Memory Search? Psychological Science, 23(7), 698–703. 10.1177/0956797612443968

Wolfe, J. M. (2021). Guided Search 6.0: An updated model of visual search. Psychonomic Bulletin & Review, 28(4), 1060–1092.

Wu, R., Pruitt, Z., Runkle, M., Scerif, G., & Aslin, R. N. (2016). A neural signature of rapid category-based target selection as a function of intra-item perceptual similarity, despite inter-item dissimilarity. *Attention, Perception*, & Psychophysics, 78(3), 749–760. 10.3758/s13414-015-1039-6

Yeh, L.-C., & Peelen, M. V. (2022). The time course of categorical and perceptual similarity effects in visual search. Journal of Experimental Psychology: Human Perception and Performance, 48(10), 1069–1082. 10.1037/xhp0001034

Yeh, L.-C., Yeh, Y.-Y., & Kuo, B.-C. (2019). Spatially Specific Attention Mechanisms Are Sensitive to Competition during Visual Search. Journal of Cognitive Neuroscience, 31(8), 1248–1259. 10.1162/jocn_a_01418

Zhang, W., & Luck, S. J. (2009). Feature-based attention modulates feedforward visual processing. Nature Neuroscience, 12(1), 24–25. 10.1038/nn.2223

